# A Fatty Acid Anabolic Pathway in Specialized-Cells Remotely Controls Oocyte Activation in *Drosophila*

**DOI:** 10.1101/2021.04.19.440456

**Authors:** Mickael Poidevin, Nicolas Mazuras, Gwénaëlle Bontonou, Pierre Delamotte, Béatrice Denis, Maëlle Devilliers, Delphine Petit, Claude Wicker-Thomas, Jacques Montagne

**Affiliations:** Institut for Integrative Biology of the Cell (I2BC), UMR 9198, CNRS, Université Paris-Sud, CEA, 91190, Gif-sur-Yvette, France; Laboratoire Evolution, Génomes, Comportements, Ecologie (EGCE), UMR 9191, CNRS, IRD, Université Paris-Sud and Université Paris-Saclay, 91190, Gif-sur-Yvette, France

**Author notes:** CWT and JM contributed equally to the work. IEES PARIS, UMR 7618, UPEC, 94010 Créteil Cedex, France. Department of Ecology and Evolution, UNIL, CH-1015 Lausanne.

## Abstract

Pheromone-mediated partner recognition is crucial for maintenance of animal species. Here, we discover a metabolic link between pheromone and gamete physiology. In female genital tract, oocyte maturation is arrested at a specific meiotic-phase. Release of this arrest, called oocyte-activation, is triggered by a species-dependent signal. We show in *Drosophila melanogaster* that oenocytes, which produce the fatty acids (FAs) used as precursors of cuticular hydrocarbons (CHCs), including pheromones, are also essential for oocyte activation. We identified a set of FA-anabolic enzymes required within oenocytes for the synthesis of a particular FA that is not a CHC precursor but controls oocyte activation. Our study thus reveals that two tightly linked FA-anabolic pathways act in parallel, one to produce sexual pheromones, the other to initiate embryonic development. Given that pheromone-deficient *Drosophila melanogaster* females are highly attractive for males irrespective of their species, this oenocyte function might have evolved to prevent hybrid development.

## INTRODUCTION

In multicellular organisms, sexual reproduction is crucial for species survival. Meeting mates requires pheromonal signals between individuals of the same species (Rekwot, Ogwu et al., 2001). After mating, sperm remains in the female genital tract, where its survival duration varies amongst species (Neubaum & Wolfner, 1999). In mammalian females, sperm entry into the oocyte triggers a calcium signal that induces oocyte activation (Kashir, Nomikos et al., 2014, Miao & Williams, 2012, Swann & Lai, 2016). The regulation of sexual reproduction integrates several physiological inputs, which are not fully characterized. Given the plethora of genetic tools allowing tissue targeted gene miss-expression, *Drosophila melanogaster* provides a convenient model system to address these issues.

In *Drosophila* females, oogenesis produces oocytes arrested at metaphase I of meiosis. Oocyte activation also depends on a calcium signal (Sartain & Wolfner, 2013), which is not triggered by sperm entry, but proceeds while the oocyte moves through the oviduct and the uterus (Heifetz, Yu et al., 2001). Activation provokes i) meiosis completion, ii) formation of the vitelline membrane that prevents polyspermy and iii) translation of maternally provided mRNAs (Aviles-Pagan & Orr-Weaver, 2018, Horner & Wolfner, 2008b, Krauchunas & Wolfner, 2013). In well-fed fertilized females, eggs are continuously produced and spermatozoa stored in the seminal receptacle and spermathecae are synchronously delivered to fertilize the oocytes, indicating that regulatory processes coordinate nutritional status with oogenesis, and egg laying with sperm delivery (Avila, Bloch Qazi et al., 2012, Wolfner, 2011).

*Drosophila* sexual pheromones are cuticular hydrocarbon (CHC) members (Montagne J. & Wicker-Thomas, 2020) that form by decarboxylation of very long chain fatty acids (VLCFAs) (Qiu, Tittiger et al., 2012). In eukaryotes, VLCFA synthesis is catalyzed by the elongase complex from a LCFA (long chain fatty acid) substrate (Jakobsson, Westerberg et al., 2006), whereas LCFA synthesis is catalyzed by FASN (fatty acid synthase) from an acetyl-CoA primer (Maier, Leibundgut et al., 2010). Synthesis of both LCFAs and VLCFAs requires the sequential incorporation of malonyl-CoA units, whose biogenesis is catalyzed by the Acetyl-CoA carboxylase (ACC) (Barber, Price et al., 2005). Regarding *Drosophila* CHCs, we previously showed that synthesis of VLCFAs takes place exclusively in the oenocytes, which are rows of cells located underneath the dorsal abdominal cuticle, whereas the LCFAs used as precursors are of flexible origin (Wicker-Thomas, Garrido et al., 2015). The *Drosophila* genome encodes three *FASN* genes, two of them, *FASN2* and *FASN3*, are specifically expressed in the oenocytes (Chung, Loehlin et al., 2014, Garrido, Rubin et al., 2015, Parvy, Napal et al., 2012). It has been shown that FASN2 catalyzes the synthesis of methylated/branched(mb)FAs, using a primer distinct from acetyl-CoA (Chung et al., 2014, Wicker-Thomas et al., 2015). Regarding FASN3, we previously reported that its knockdown affects tracheal waterproofing in larvae (Parvy et al., 2012) and desiccation resistance but not CHC synthesis in adult flies (Wicker-Thomas et al., 2015). Here, we show that a particular FA-anabolic pathway operating from the oenocytes remotely controls oocyte activation. Our study thus suggests a metabolic link between pheromone synthesis and female fertility.

## RESULTS

### Identification of FA-metabolic genes required in the oenocytes for female fertility

While studying CHCs, we observed that *Drosophila* females deficient for FASN3 in their oenocytes exhibited a sterile phenotype. To identify additional genes potentially involved in this process, we directed inducible interfering RNA (*RNAi*) to 57 genes encoding enzymes related to FA-metabolism, using the *1407-gal4* driver that is active in oenocytes from mid L3 larval stage to adulthood (Wicker-Thomas et al., 2015). In their progeny, at least twenty virgin females—hereafter referred as *1407>geneX-RNAi*—were individually crossed to Canton-S males. Next, females were transferred every second day to new vials and the number of emerging adults was counted. This way, we identified four additional gene products required for fertility (Table EV1 and Fig EV1A), including ACC, a component of the elongase complex (KAR), a bipartite FA-transporter/acyl-CoA ligase (FATP) and CG6432, which encodes a putative short chain acyl-CoA ligase. Sibling males of the sterile females were tested for their ability to fertilize Canton-S females, yet none of them appeared to be sterile (Fig EV1B). Taken together, these findings reveal that a VLCFA produced in the oenocytes controls female but not male fertility.

### Searching for oenocyte defects

*KAR*, which encodes a component of the elongase complex, has also been shown to prevent oenocyte degeneration in adult flies (Chiang, Tan et al., 2016). We therefore, investigated whether the oenocyte knockdowns that produced female sterility also affected oenocyte viability in females. Consistently, *1407>KAR-RNAi* resulted in oenocyte degeneration in 27-day old females, but not in 10-and 18-day old females (Fig EV2A-F). As previously described (Wicker-Thomas et al., 2015), *1407>ACC-RNAi* induced lipid accumulation in the oenocytes but did not affect their viability (Fig EV2G-I), whereas *1407>FATP-RNAi* resulted in oenocyte degeneration that was visible as of day 18 (Fig EV2J-L). In contrast, *1407>FASN3-RNAi* and *1407>CG6432-RNAi* flies were fully viable and the oenocytes of females did not degenerate, did not accumulate high amounts of lipid droplets and appeared similar to controls (Fig EV2M-R). Following knockdown of any of the genes identified, the female sterile phenotype was invariably observed as of day 8, ie. when oenocytes appeared viable (Fig EV2D,G,J,M,P), indicating that the sterile phenotype resulted from the inactivation of a specific FA metabolic pathway, rather than deficient oenocytes.

### Metabolic pathway required for female fertility

*CG6432* is proposed to encode a short chain acyl-CoA ligase (Flybase, 2003). However, protein blast analysis revealed that it contains a putative propionyl-CoA synthase domain and that its closest homologue is a short chain FA-acyl-CoA ligase in mouse (Acss3), and an acetyl-CoA ligase in yeast (Acs1) (Fig EV3). Therefore, we investigated whether *CG6432* was also required in other FA-anabolic pathways. We previously reported that oenocyte knockdown of *ACC, FASN3, KAR* and *FATP* in young larvae results in flooding of the tracheal system and in lethality at the L2/L3 transition (Parvy et al., 2012). As previously reported for FASN3 (Garrido et al., 2015), this phenotype also happened for oenocyte or ubiquitous knockdown of *CG6432* (Fig 1A), indicating that the CG6432 gene product resides in the oenocyte-specific metabolic pathway that controls tracheal watertightness in larvae. We also previously reported that knockdown of *ACC, KAR* and *FATP*, but not of *FASN3*, in adult oenocytes results in a drop of total CHC amounts (Wicker-Thomas et al., 2015). Knockdown of *CG6432* in adult oenocytes did not reduce total CHC amounts but resulted in a dramatic drop of methylated/branched(mb)CHCs (Table EV2, Fig 1B and Fig EV4), a phenotype previously reported in *FASN2* oenocyte-knockdown flies (Chung et al., 2014, Wicker-Thomas et al., 2015). In contrast to *FASN1*, fat body knockdown of *CG6432* did not reduce total triacylglycerol levels (Fig 1C), indicating that it is not required for FASN1 activity in the fat body. Taken together, these findings reveal that the enzyme encoded by *CG6432* selectively resides in the FASN2 and FASN3 but not FASN1 anabolic pathway.

**Figure 1:**
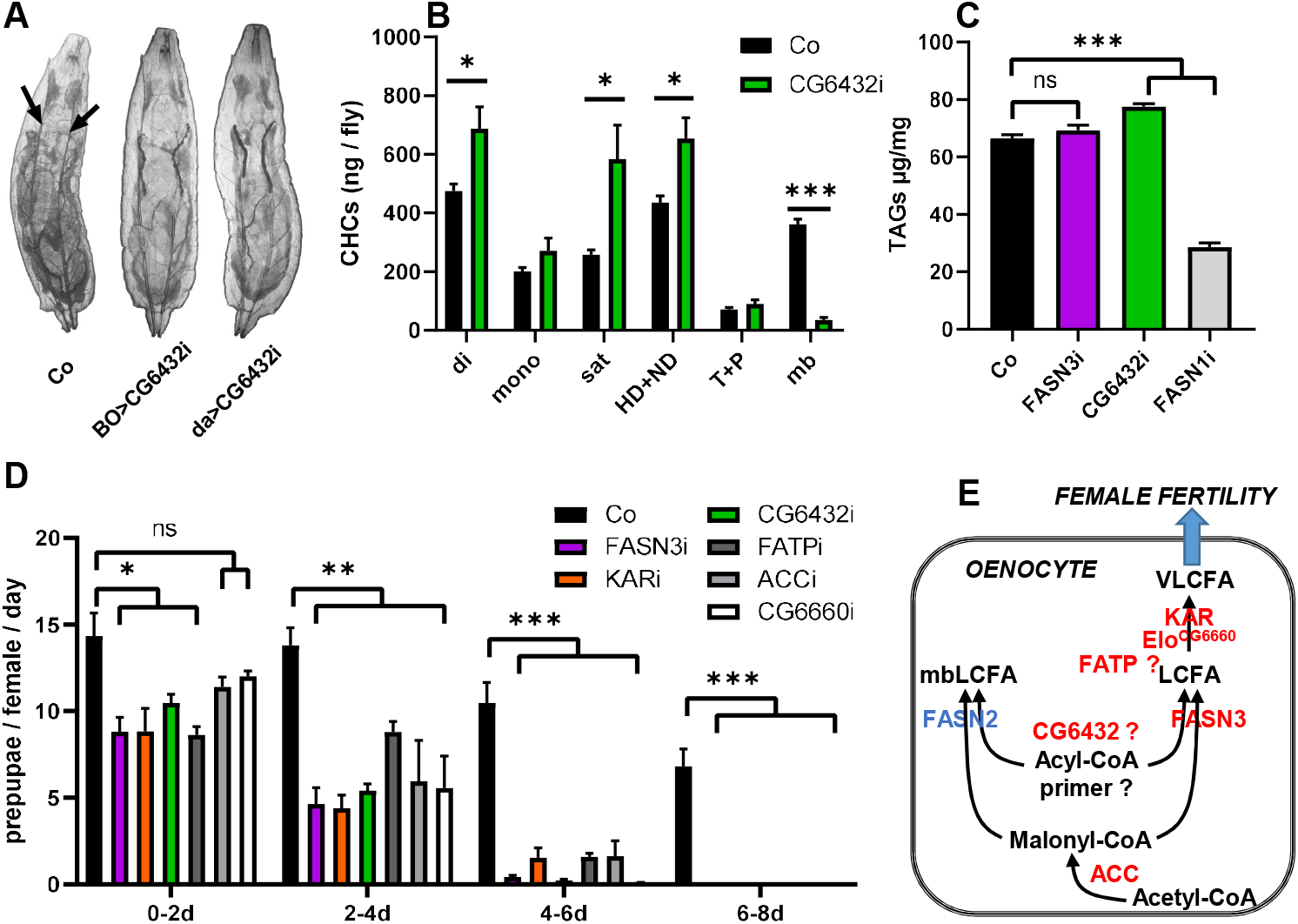
Characterization of an oenocyte FA anabolic pathway required for female fertility: (**A**) Oenocyte(BO)-or ubiquitous(da)-knockdown of *CG6432* induced tracheal flooding in late L2 larvae; in their flooded section, the tracheal trunks (arrows in control: Co) were hardly visible. (**B**) CHCs amounts in control (black) or *1407>6432-RNAi* (green) females (n=10); Amounts of dienes (di), monoenes (mono), saturated linear (sat), pheromones (HD+ND) and mbCHCs (mb) are listed in Table EV2; note the strong reduction of mbCHCs. (**C**) TAG content in 0-5 hours prepupae of the following genotypes: *Cg-gal4* control (black), *Cg>FASN3-RNAi* (purple), *Cg>CG6432-RNAi* (green) and *Cg>FASN1-RNAi* (grey). (**D**) Pupal progeny of *promE-gal4* females either control (black) or expressing an RNAi to *FASN3* (purple), *KAR* (orange), *CG6432* (green), *FATP* (dark grey), *ACC* (light grey) or *elo*^*CG6660*^ (white). Five females were mated with five males for two days (0-2d), then, males were removed and females transferred in new tubes every second day (2-4d, 4-6d, 6-8d). Each bar represents the man value of 3-5 replicates. (**E**) Oenocyte anabolic pathway producing a VLCFA controlling fertility, where CG6432 and FATP are potential acyl-CoA synthase for the primer used by FASN2/FASN3 and for FA elongation, respectively; enzymes (blue/red) and metabolites (black) are indicated.

Next, we made use of the *promE-gal4* driver, which, in contrast to the *1407-gal4* driver (Wicker-Thomas et al., 2015), is active only in the oenocytes in *Drosophila* females as of stage L1 (Billeter, Atallah et al., 2009). All the genes required in adult oenocytes for female fertility (Table EV1) are also essential in larval oenocytes for spiracle watertightness, except the elongase subunit *elo*^*CG6660*^, the knockdown of which seemed to affect only the latter process (Parvy et al., 2012). Therefore, to retest *elo*^*CG6660*^ with the *promE-gal4* driver we used a different RNAi line (Table EV1). Given that *promE-gal4* induced larval lethality when driving any of the genes of interest, its activity was blocked until early metamorphosis using the thermo-sensitive Gal4 inhibitor, Gal80^ts^. RNAi expression was induced by a temperature shift to 27°C. Emerging females were maintained at 27°C, mated 3-5 days after virgin collection and changed to new vials every second day (Fig 1D). In this way, newly fertilized females expressing any of the RNAis of interest exhibited a net reduction of fertility compared to that of control females, which quickly dropped to complete sterility (Fig 1D). Moreover, in contrast to the fertility of *1407>elo*^*CG6660*^*-RNAi* females (Fig EV1A), *promE-gal4*>*elo*^*CG6660*^*-RNAi* females appeared to be sterile, possibly because of different RNAi strength (Fig 1D). Taken together, these findings reveal that a FA anabolic pathway working in the oenocytes produces a particular VLCFA that is required for female fertility (Fig 1E). While the acyl-CoA primer for mbLCFA synthesis catalyzed by FASN2 is likely a propionyl-CoA, whereas the one used by FASN3 is yet unknown. Thus, it is tempting to speculate that CG6432 is responsible for the synthesis of this unconventional FA primer (Fig 1E).

### Oenocytes control female fertility

To get further insights into this oenocyte function, we focused on *FASN3* and *CG6432*, since both these genes are essential only in the oenocytes and their knockdown alters neither oenocyte viability (Fig EV2) nor CHC synthesis (Fig 1B and (Wicker-Thomas et al., 2015)). Furthermore, confocal imaging of oenocytes (Fig 2A) and ovaries (Fig 2B) of *promE>FASN3-RNAi* and *promE>CG6432-RNAi* females dissected 10 days after mating, revealed no apparent defects, all the egg chamber stages were visible (Fig 2B). Moreover, in mating assays, single wild type males did not exhibit any significant preference when given a choice between a wild type female and a female of either genotype *promE>FASN3-RNAi* or *promE>CG6432-RNAi* (Fig 2C). Next, we monitored the number of eggs laid by females. Control, *promE>FASN3-RNAi* and *promE>CG6432-RNAi* females laid high amounts of eggs the day after fertilization (Fig 2D); this number decreased the following days, although this effect was more pronounced for *promE>FASN3-RNAi* and *promE>CG6432-RNAi* compared to control females (Fig 2D). However, consistent with a fertility defect, the number of eggs that formed pupae in the *FASN3-*and *CG6432-RNAi* conditions dropped more dramatically than the number of eggs laid, as compared to control (Fig 2E-F). Taken together, these findings reveal that the oenocyte metabolic pathway that controls female fertility affects neither partner mating nor oogenesis.

**Figure 2:**
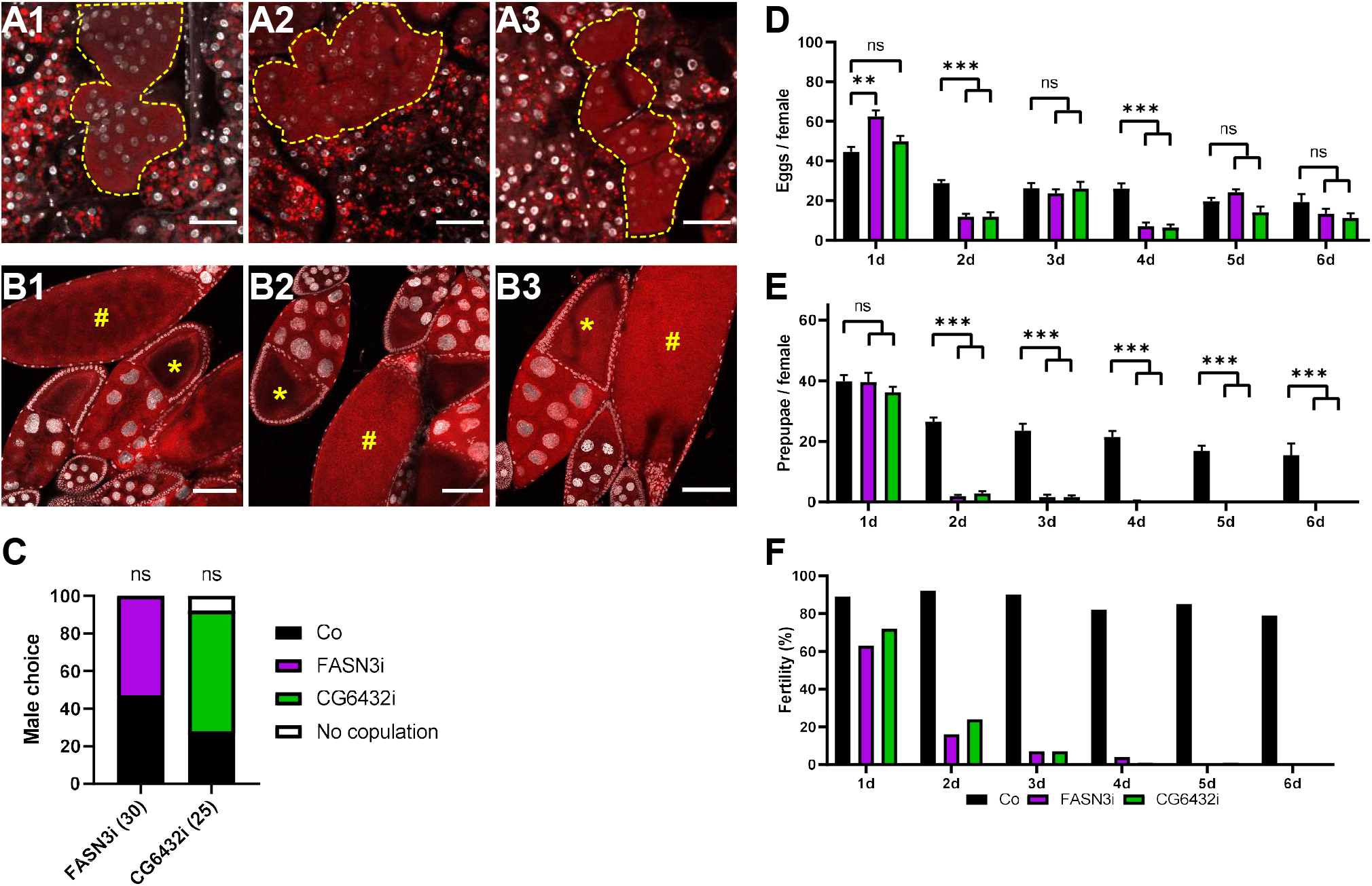
Oogenesis and mating: (**A-B**) Oenocytes (yellow dotted line in A1-3) and stage 9-10 (**\***) and late (**#**) egg chambers (B1-3) of *promE-gal4* females either control (A1, B1), or directing an RNAi to *FASN3* (A2, B2) or *CG6432* (A3, B3); tissues were dissected 10-days after mating; lipids and nuclei were labelled with Nile Red (red) and DAPI (silver), respectively; scale bars: 40 µm (A1-3) and 100 µm (B1-3). (**C**) Mating choice of single wild type males in the presence of two females, one control and one expressing an RNAi to either *FASN3* (left) or *CG6432* (right); bars represent the percentage of copulation with control (color) or RNAi-expressing (purple or green) females; males tend to prefer *CG6432-RNAi* females although not significantly. (**D-F**) eggs (D) and pupae (E) from *promE-gal4* females either control (black) or expressing an RNAi to *FASN3* (purple) or *CG6432* (green); three females were mated with three males for one day, then, males were removed and females transferred in new tubes every day over a 6-day period; index of fertility (F) were evaluated as the ratio of prepupae to eggs.

### Oenocytes do not control sperm delivery to the oocytes

After copulation, spermatozoa are stored for several days within the seminal receptacle and spermathecae of *Drosophila* females (Wolfner, 2011). We searched for potential defects in sperm storage or delivery. In control females, the number of spermatozoa in the seminal receptacle progressively decreased after mating, while females laid eggs continuously (Fig 3A); *promE>FASN3-RNAi* and *promE>CG6432-RNAi* females contained roughly the same number of spermatozoa in their seminal receptacle the day after mating, but surprisingly, this number decreased much less during the following days compared to control females (Fig 3A). Next, we monitored sperm motility in the seminal receptacle but did not observe a difference in sperm speed in *promE>FASN3-RNAi* and *promE>CG6432-RNAi* females compared to control females (Fig 3B and movies EV1-3). Moreover, confocal analysis revealed that the eggs of control, *promE>FASN3-RNAi* and *promE>CG6432-RNAi* females were fertilized as shown by the presence of sperm flagella (Fig 3C-E). These findings indicate that, despite sperm retention in storage organs, the defect of fertility is not due to a failure of sperm entry into the oocyte.

**Figure 3:**
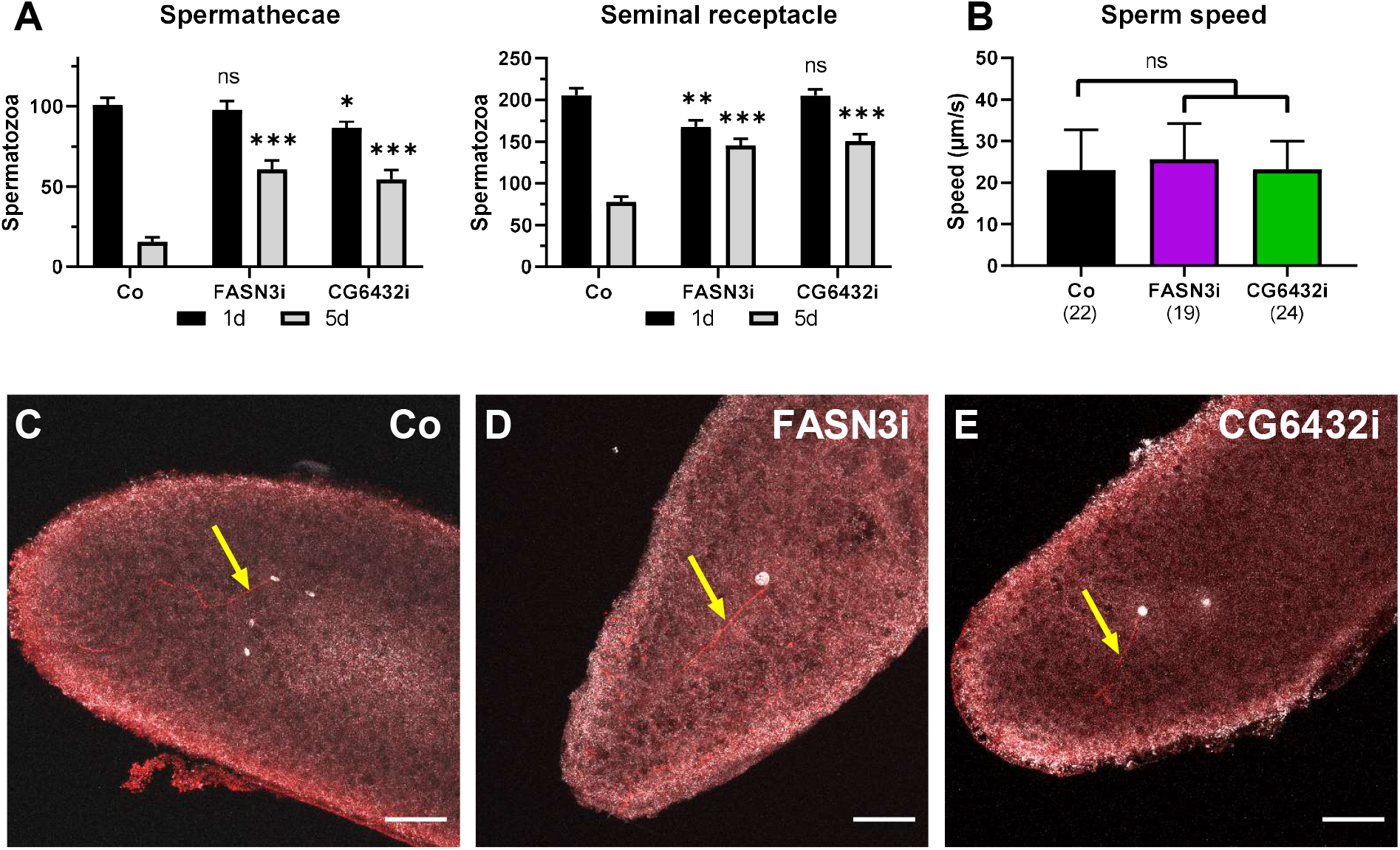
Sperm activity: (**A**) Sperm numbers in the spermathecae (left) and seminal receptacle (right) of *promE-gal4* females either control (Co) or directing an RNAi to *FASN3* or *CG6432*, one day (black) or five days (grey) after mating (n>25). (**B**) Movement speed of spermatozoa in the seminal receptacle of *promE-gal4* females either control (black), or directing an RNAi to *FASN3* (purple) or *CG6432* (green). (**C**-**E**) Eggs laid from *promE-gal4* fertilized females either control (C), or directing an RNAi to *FASN3* (D) or *CG6432* (E); eggs were collected for 40 mn, nuclei were labeled with DAPI (silver) and sperm flagella (arrows) with an anti-acetylated-tubulin (red); scale bars: 40 µm.

### Oenocytes control oocyte activation

A fertilized egg contains potentially five nuclei, the sperm and oocyte pronuclei, plus three polar globules. However, while analyzing the presence of spermatozoa, we noticed an apparent defect in the number of nuclei in *promE>FASN3-RNAi* and *promE>CG6432-RNAi* females (Fig 3C-E). We therefore, counted the number of nuclei in eggs laid by virgin females and observed that the number of nuclei in control eggs varid from zero to four, likely because some nuclei are not stained or not visible (Fig 4A and 4D). Nevertheless, the number of nuclei in *promE>FASN3-RNAi* and *promE>CG6432-RNAi* females was much lower than in control females (Fig 4B-C and 4D). Production of the three polar globules results from meiosis completion (Page & Orr-Weaver, 1997), suggesting that this process does not fully operate in *promE>FASN3-RNAi* and *promE>CG6432-RNAi* females. Meiosis completion is triggered by oocyte activation, which also induces formation of the vitelline membrane and translation of maternally provided mRNAs (Horner & Wolfner, 2008a). The vitelline membrane allows resistance to bleach induced egg lysis (Horner, Czank et al., 2006). Importantly, we observed that fertilized eggs laid by *promE>FASN3-RNAi* and *promE>CG6432-RNAi* females were much less resistant to bleach treatment (Fig 4E). However, the bleach concentration required for egg lysis was higher than the one described by others (Horner et al., 2006). Finally, we analyzed Smaug, a protein encoded by a maternally provided mRNA, whose translation is induced by egg activation (Horner & Wolfner, 2008a, Tadros, Goldman et al., 2007). Western-blot analysis revealed that Smaug was present at much lower levels in eggs laid by *promE>FASN3-RNAi* and *promE>CG6432-RNAi* than in eggs laid by control females (Fig 4F). Taken together these findings reveal that a particular VLCFA produced in the oenocytes controls oocyte activation.

**Figure 4:**
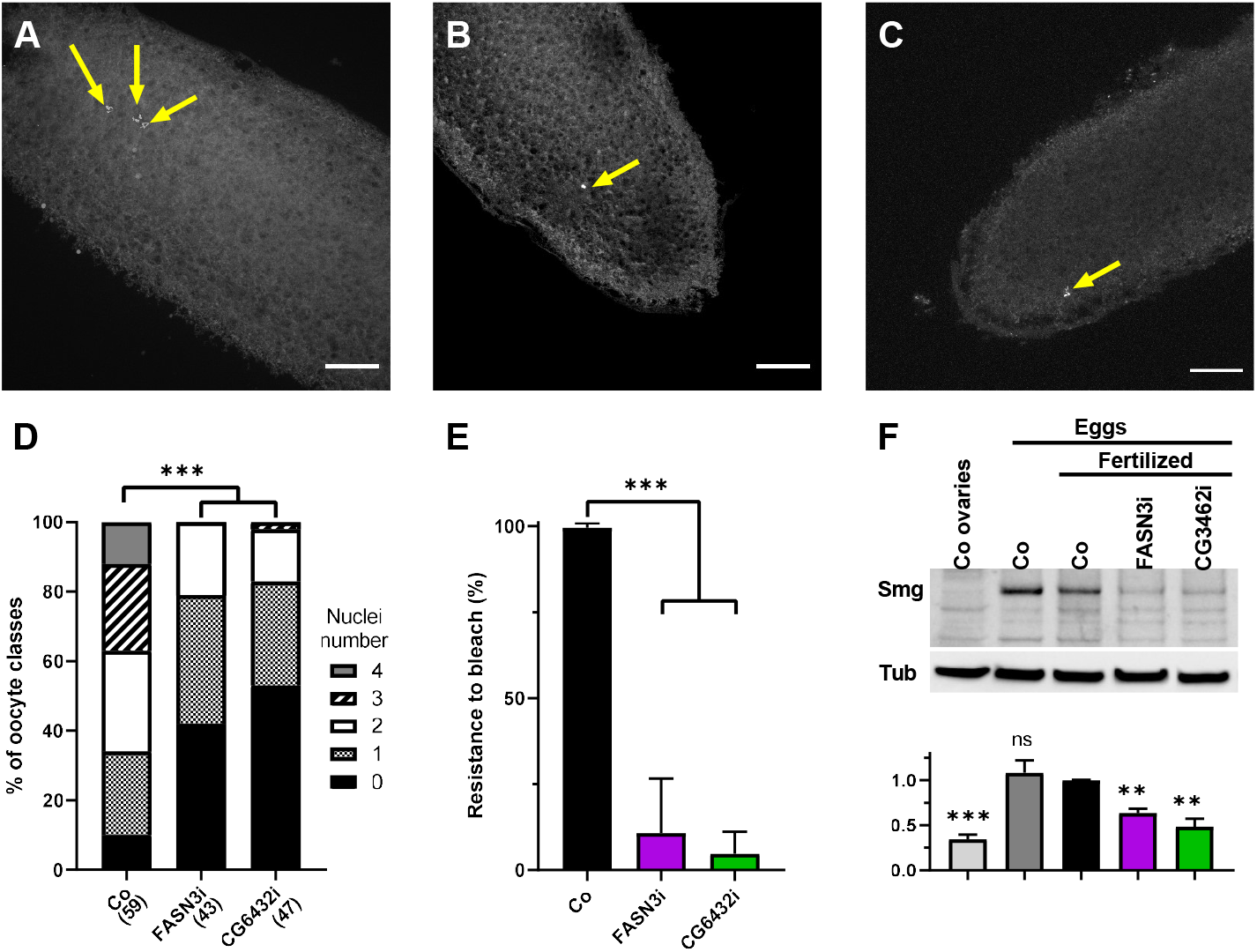
Oocyte activation: (**A**-**C**) Eggs laid from *promE-gal4* virgin females either control (A), or directing an RNAi to *FASN3* (B) or *CG6432* (C); eggs were collected for 2 hrs and nuclei were labeled with DAPI (silver); scale bars: 40 µm. (**D**) Number of nuclei counted in the egg collections (A-C). (**E**) Resistance to bleach lysis of eggs laid by *promE-gal4* fertilized females either control (black), or directing and RNAi to *FASN3* (purple) or *CG6432* (green); bars represent means from 27 independent tests, each containing 25 eggs per genotype. (**F**) Western-blotting to Smaug from protein extracts of dissected ovaries or eggs laid by *promE-gal4* females either virgin or fertilized; ovaries and unfertilized eggs were from control females, fertilized eggs were from females either control or directing an RNAi to *FASN3* or *CG6432*. The graph at the bottom compares the means of four independent blots, where the band intensity of Smaug was normalized to that of the tubulin loading control.

## DISCUSSION

In this study, we provide evidence that a FA-anabolic pathway, which takes place within the oenocytes of *Drosophila* females, remotely controls fertility. We have identified six FA-anabolic enzymes required for this process: ACC, FASN3, CG6432, FATP, KAR and Elo^CG6660^. ACC, the rate-limiting enzyme for FA synthesis, catalyzes malonyl-CoA synthesis (Barber et al., 2005). FASN3 is one of the three *Drosophila* FASN enzymes (Garrido et al., 2015). CG6432 that encodes a putative short chain acyl-CoA synthase, might catalyze the synthesis of the acyl-CoA primer used by FASN2 and FASN3 to build up FAs (Fig 1D). FATP is a bipartite fatty acid transporter/acyl-CoA synthase, although our previous study supports that it works instead as an acyl-CoA synthase tightly linked to VLCFA synthesis (Wicker-Thomas et al., 2015). KAR and Elo^CG6660^ are subunits of the elongase complex that comprises two reductases (KAR and TER), a dehydratase and the elongase (Jakobsson et al., 2006). The elongase subunit assigns specificity to FA primer usage and to the VLCFA produced; both of which remain unidentified for Elo^CG6660^, as for most of the 20 elongases encoded by the *Drosophila* genome (Montagne J. & Wicker-Thomas, 2020). Nonetheless, the oenocyte-mediated control of female fertility likely relies on either a single or a restricted subclass of VLCFA(s), which might be the precursor of a yet unknown lipid hormone. We also observed that oenocyte knockdown of *FATP* and *KAR*, but not of *ACC, FASN3* and *CG6432*, results in oenocyte degeneration, further supporting the notion that FATP operates closely to VLCFA synthesis (Wicker-Thomas et al., 2015). Potentially, the degeneration process is not induced by the lack of VLCFAs but by the accumulation of LCFA precursors, which happens when inhibiting VLCFA but not LCFA synthesis. These six enzymes are also required in larval oenocytes to maintain the spiracle watertightness (Parvy et al., 2012), suggesting that FASN3-dependent VLCFAs are the precursors of lipid messengers transmitted through the haemolymph to their target tissues. Our study demonstrates that one of these particular VLCFAs operates on oocyte activation. In females deficient for this VLCFA, i) the low number of nuclei in unfertilized eggs suggests that meiosis completion does not fully operate, ii) the bleach sensitivity of eggs argues for a vitelline membrane defect, and iii) the low levels of Smg suggests a defect in the translation of maternally provided mRNAs. All these processes are typically triggered by oocyte activation (Aviles-Pagan & Orr-Weaver, 2018). Nonetheless, the lack of this remote control is unlikely to result in a total blockade of oocyte activation, since all these processes are affected but not fully suppressed.

At first glance, it is surprising that a VLCFA synthesized through a metabolic pathway tightly parallel to the one responsible for pheromone biogenesis, remotely controls oocyte activation. Of note, it has been shown that *Drosophila melanogaster* females devoid of oenocytes are attractive to males from other species, including *D. simulans, D. yakuba* and *D. erecta (Billeter et al*., *2009)*. These females are also more attractive to wild type *Drosophila melanogaster* males, since the delay for copulation with oenocyte deficient females in shortened. The increased attractiveness likely depends on carboxyl-methylated FAs produced by *Drosophila* females, since shortening of the copulation delay does not happen with males deficient for the odorant receptor of these carboxyl-methylated FAs. These studies support the notion that female attractiveness depends on a balance between repulsion and attraction, where the cocktail of CHC-related compounds would be rather repulsive, while the species distinctive pheromone signature confers the selective attractiveness for conspecific males. Given that pheromone biogenesis shares common enzymes with the synthesis of the oocyte-activating VLCFA, deficiency in the former pathway should also impede the latter. Therefore, it is tempting to speculate that these two pathways have been co-selected throughout evolution to prevent hybrid development. Flies harboring non-functional oenocytes would copulate with males irrespective of their species with the resulting hybrids being potential competitors. Identification of the FASN3-dependent VLFCAs will constitute the next challenge and should allow deciphering its action mechanism on oocyte activation and determining whether a similar regulatory process is conserved throughout evolution to favor species isolation.

## Acknowledgements

We wish to thank Benjamin Loppin for critical advices, VDRC, NIG-FLY M Simonelig for fly stocks and reagents, and Melanie Gettings for manuscript editing. Thanks are due to funding supports from Centre National de la Recherche Scientifique (to CWT, JM), IFR115 grant (to CWT and JM), *Fondation ARC* (1555286 to JM), French league against Cancer (M27218 to JM), French Government (fellowships MENRT 2015-155 to MD and 2020-110 to PD).

## Author contributions

CWT and JM conceived the study; MP, NM, PD, BD, MD, DP, CWT and JM performed the methodology; MP, GW, CWT and JM analyzed the data; JM wrote the manuscript. All authors reviewed the manuscript.

## MATERIAL AND METHODS

### Genetics and fly handling

#### Fly stock

*1407-gal4* (oenocytes from mid L3 stage) (Ferveur, Savarit et al., 1997), *BO-gal4* (oenocytes in embryo and early larvae) (Gutierrez, Wiggins et al., 2007), *promE-gal4* (oenocytes from L1 stage) (Billeter et al., 2009), *P[w8, ProtB-DsRed-monomer, w+]50A UAS-GFP* (Manier, Belote et al., 2010), *Cg-gal4* (fat body), *da-gal4* (ubiquitous), *Tub-gal80ts* (ubiquitous) from BDSC (https://bdsc.indiana.edu). Inducible *UAS-RNAi* lines (Table S1) from VDRC (https://stockcenter.vdrc.at/control/main) (Dietzl, Chen et al., 2007), NIG (https://shigen.nig.ac.jp/fly/nigfly) or previously described (Palm, Sampaio et al., 2012, Parvy et al., 2012)

### Mating and related analyses

Mating choice were performed as previously described (Wicker-Thomas et al., 2015). For sperm counting and speed, females were fertilized with *P[w8, ProtB-DsRed, w+]* males that labelled sperm nuclei. For counting, spermathecae and seminal receptacle were dissected in PBS 1X, fixed 20 min in PFA 4%, mounted in DABCO and spermatozoa were counted with a Zeiss Imager M2 fluorescent microscope. For sperm speed, seminal receptacle were dissected and mounted in Biggers, Whitten and Whittingham modified medium (95 mM NaCl, 4.8 mM KCl, 1.3 mM CaCl2, 1.2 mM MgSO4, 1.2 mM KH2PO4, 5.6 mM glucose, 25 mM NaHCO3, 20mM HEPES, 0.6% fatty acid free BSA, pH 7.6), supplemented with 0.5 mM trehalose. Time lapse were imaged with Zeiss Imager M2 fluorescent microscope (10 seconds; 0.15 second frame interval); speed means are from at least 20 spermatozoa per genotype (five females each). For bleach resistance, eggs were collected for 2 hrs, incubated with commercial bleach (3,7% chlorax) for 10 mn, rinsed with water, and numbers of eggs still visible were counted.

### Imaging

Dissected oenocytes and ovaries were fixed and labeled with DAPI and Oil-Red-O as previously described (Wicker-Thomas et al., 2015). For sperm flagella immunostaining, eggs were dechorionated for 2 min in commercial bleach (3,7% chlorax), fixed in a 1:1 heptane:methanol mixture and stored at −20°C. Next, embryos were washed three times for 10 min with PBS 0.1% Triton X100, incubated with primary antibody (anti-acetylated tubulin; Sigma-Aldrich; T6793) and DAPI on a rotating wheel overnight at 4°C, washed three times (20 min each) and incubated with an anti-mouse antibody Alexa Fluor 568 nm (Invitrogen; A11061). Samples were mounted in DAPCO and analyzed on a Leica SP8 confocal laser-scanning microscope.

### Biochemistry

Experiments performed as previously described: CHC measurement (Wicker-Thomas et al., 2015). TAG measurement (Garrido et al., 2015); protein extracts and western-blotting (Montagne, Lecerf et al., 2010). The Smaug antibody was kindly provided by M Simonelig (Chartier, Klein et al., 2015). Quantification of western-blot was performed using ImageJ.

### Statistics

Statistical analysis were performed using PRISM/Graphpad. Statistical significance were indicated as *, ** and *** corresponding to P< 0.05, 0.01 and 0.001, respectively. T-test were used for Fig. 1B, 1C, 1D, 2D, 2E, 3A, 3B, 4E, 4F. Chi-2 test were used for Fig. 2C and 4D. ANOVA was used for Table S2.

## EXTENDED VIEW

Movie EV1. Displacement of spermatozoa in the seminal receptacle of *promE-gal4* control females.

Movie EV2. Displacement of spermatozoa in the seminal receptacle of *promE-gal4*>*FASN3-Ri*.

Movie EV3. Displacement of spermatozoa in the seminal receptacle of *promE-gal4*>*CG6432-Ri*.

**Figure EV1:**
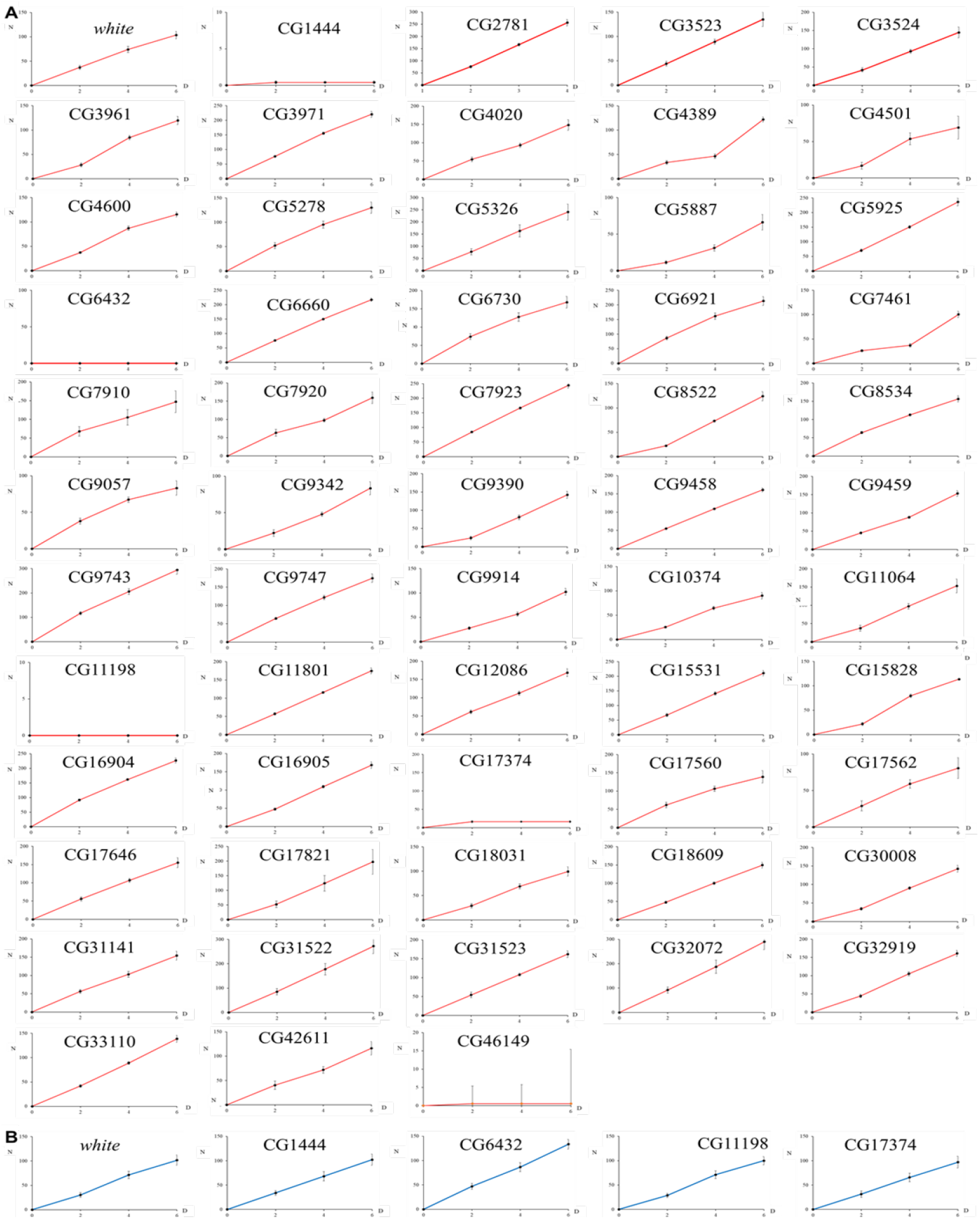
Screening for sterility. (**A**) *1407-gal4>UAS-RNAi* females crossed to Canto-S males were let to lay eggs during six days (D) in three successive vials and the progeny was counted at adult emergence (N). (**B**) Reciprocal crosses to test male fertility.

**Figure EV2:**
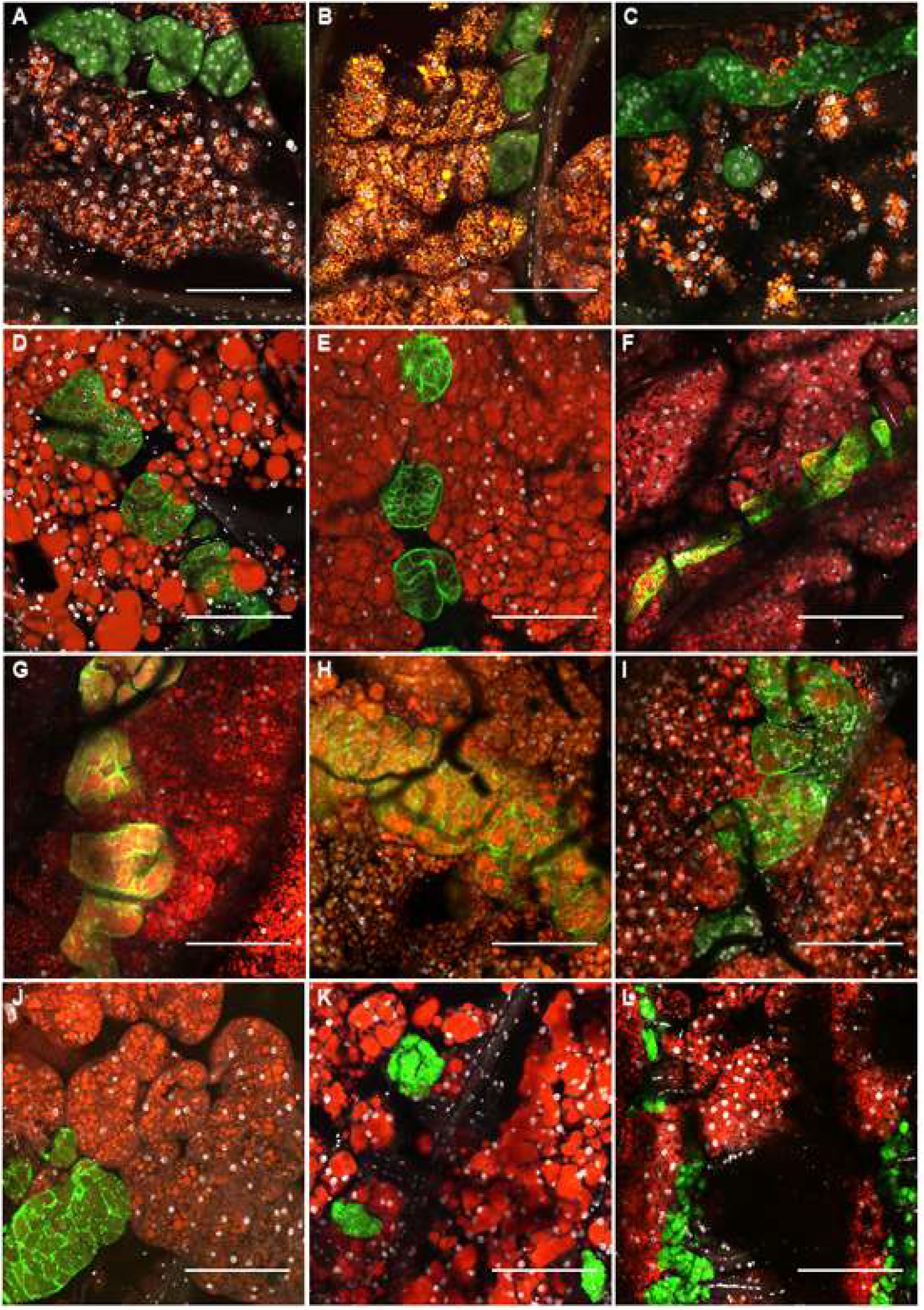
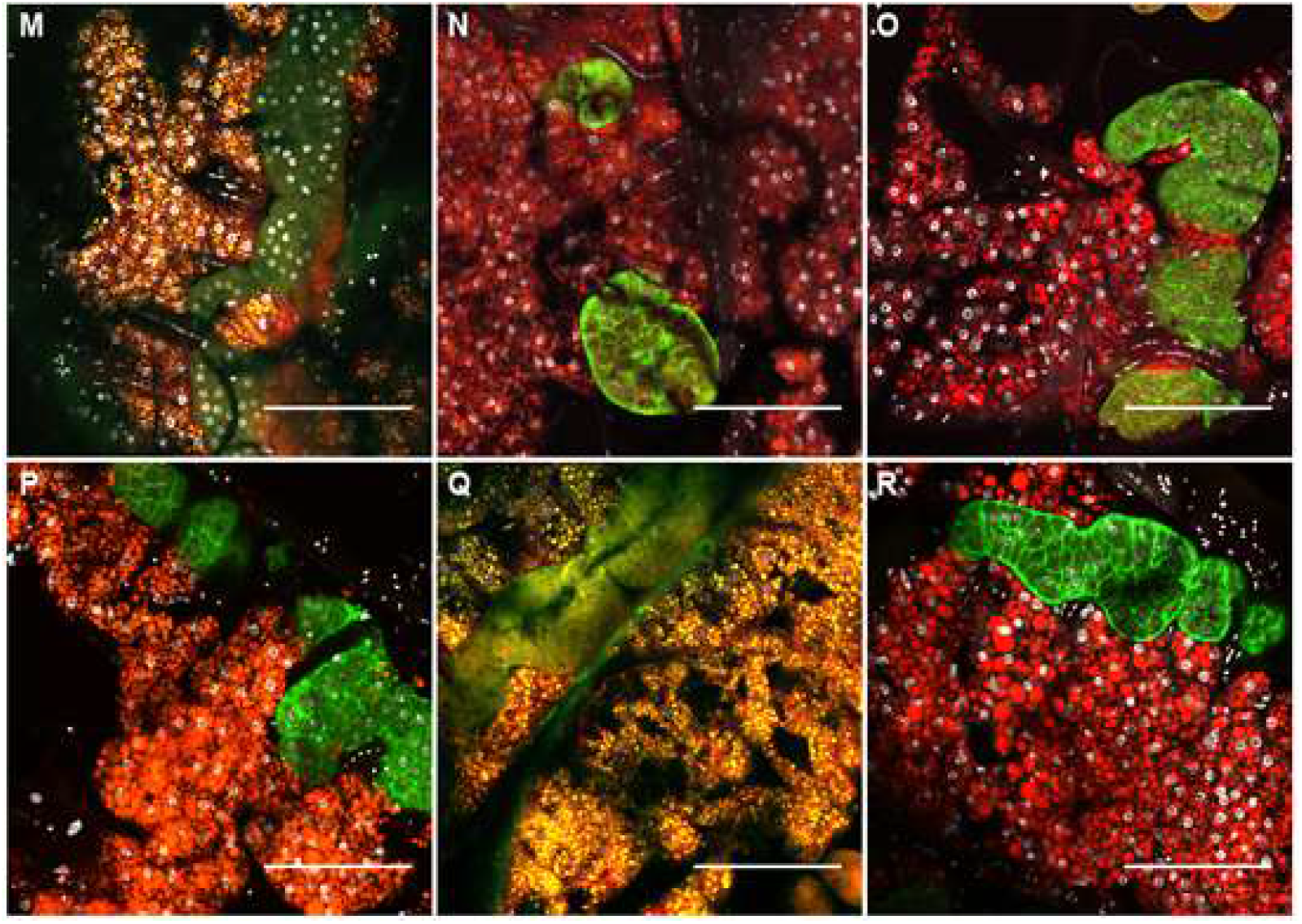
Oenocytes structure during lifespan. Dorsal abdominal cuticles stained for lipids and nuclei of *1407>UAS-GFP* females either control (A-C), or directing UAS-RNAis to KAR (D-F), ACC (G-I), FATP (J-L), FASN3 (M-O) or CG6432 (P-R). Females were dissected 9 days (A,D,G,J,M,P), 18 days (B,E,H,K,N,Q) or 27 days (C,F,O,R) after adult emergence, except ACC-and FATP-RNAis flies that did not survive longer than 24 days (I,L). Oenocytes were visualized by GFP (green) the nuclei by DAPI (silver) and the fat body by Nile red. Since the Nile red partially interferes with the GFP channel and that the GFP intensity varied a lot for unknown reasons, the lipid staining appeared either red (strong GFP) or orange (low GFP). Note that oenocyte loss appeared earlier in age for FATP-RNAi (K,L) than for KAR-RNAi (F). Scale bars: 100μm.

**Figure EV3:**
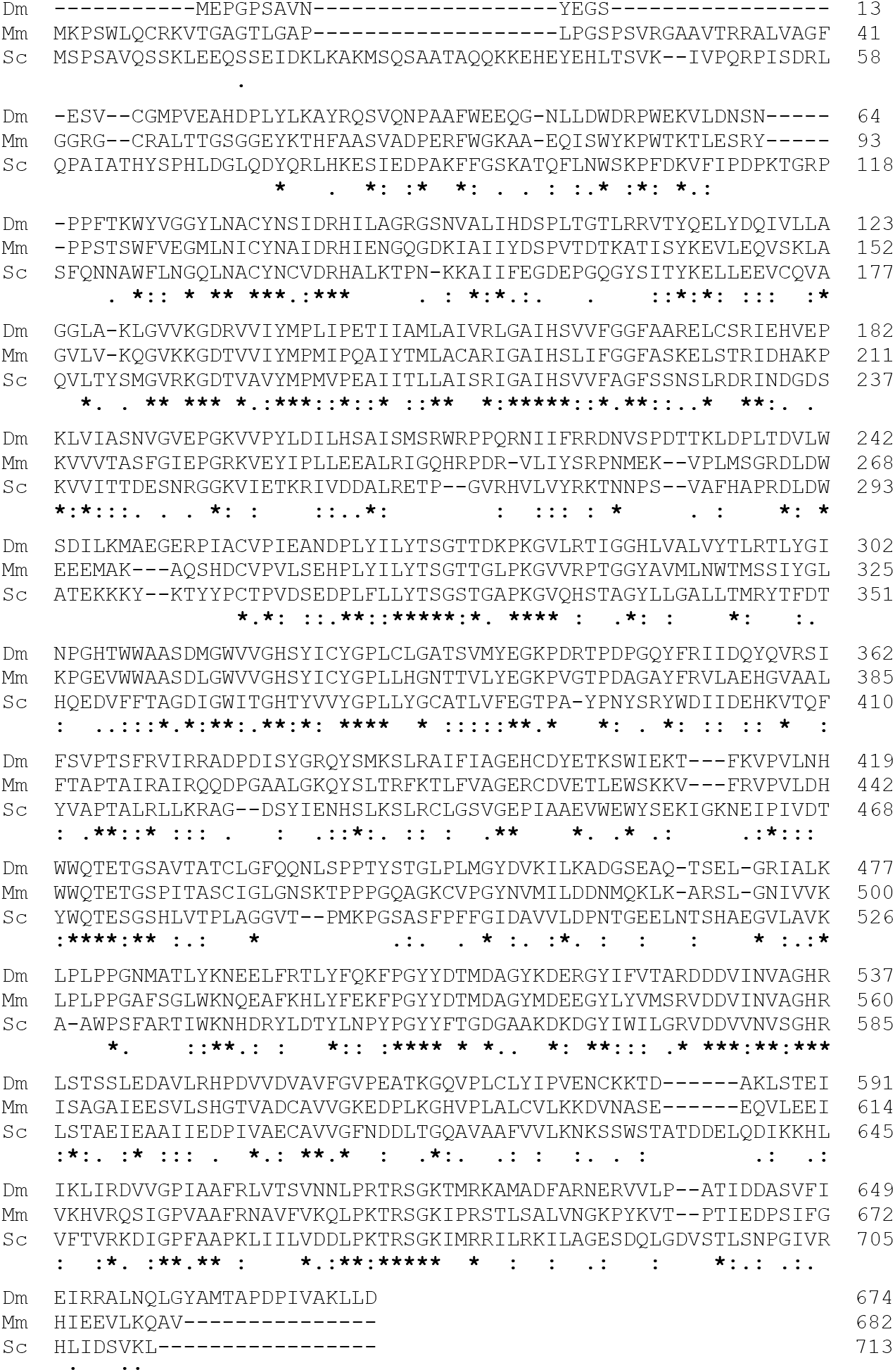
CG6432 blast. Peptide sequence alignment of CG6432 (Dm) to the best homologues Acss3 in mouse (Mm) and Acs1 in yeast (Sc), using www.uniprot.org.

**Figure EV4:**
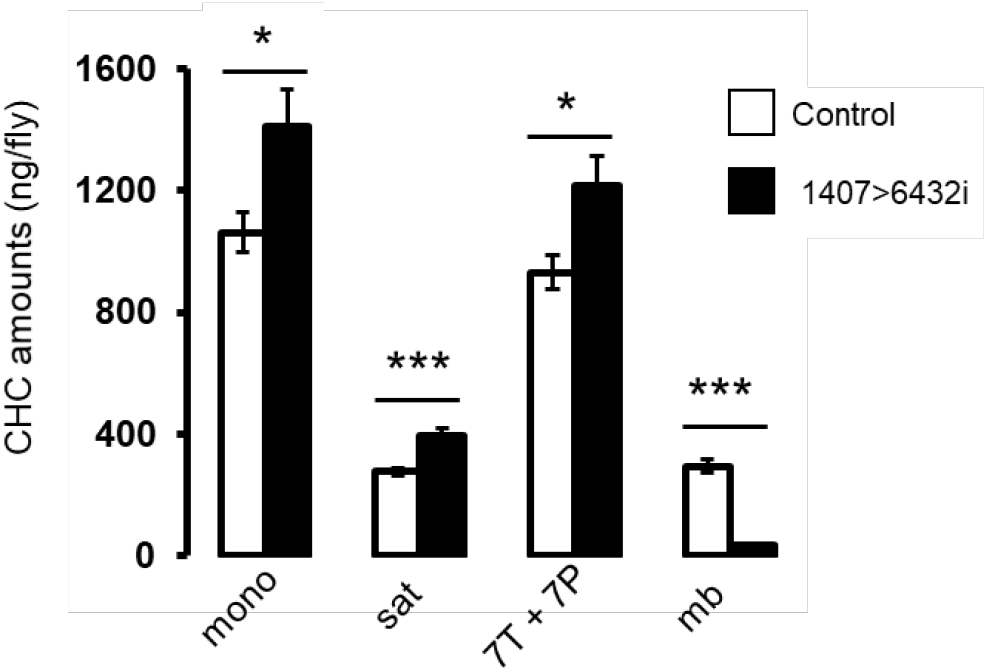
male CHCs. Means values of CHCs from 10 control (white) or 10 *CG6432-RNAi* (black) males.

**Table EV1.**
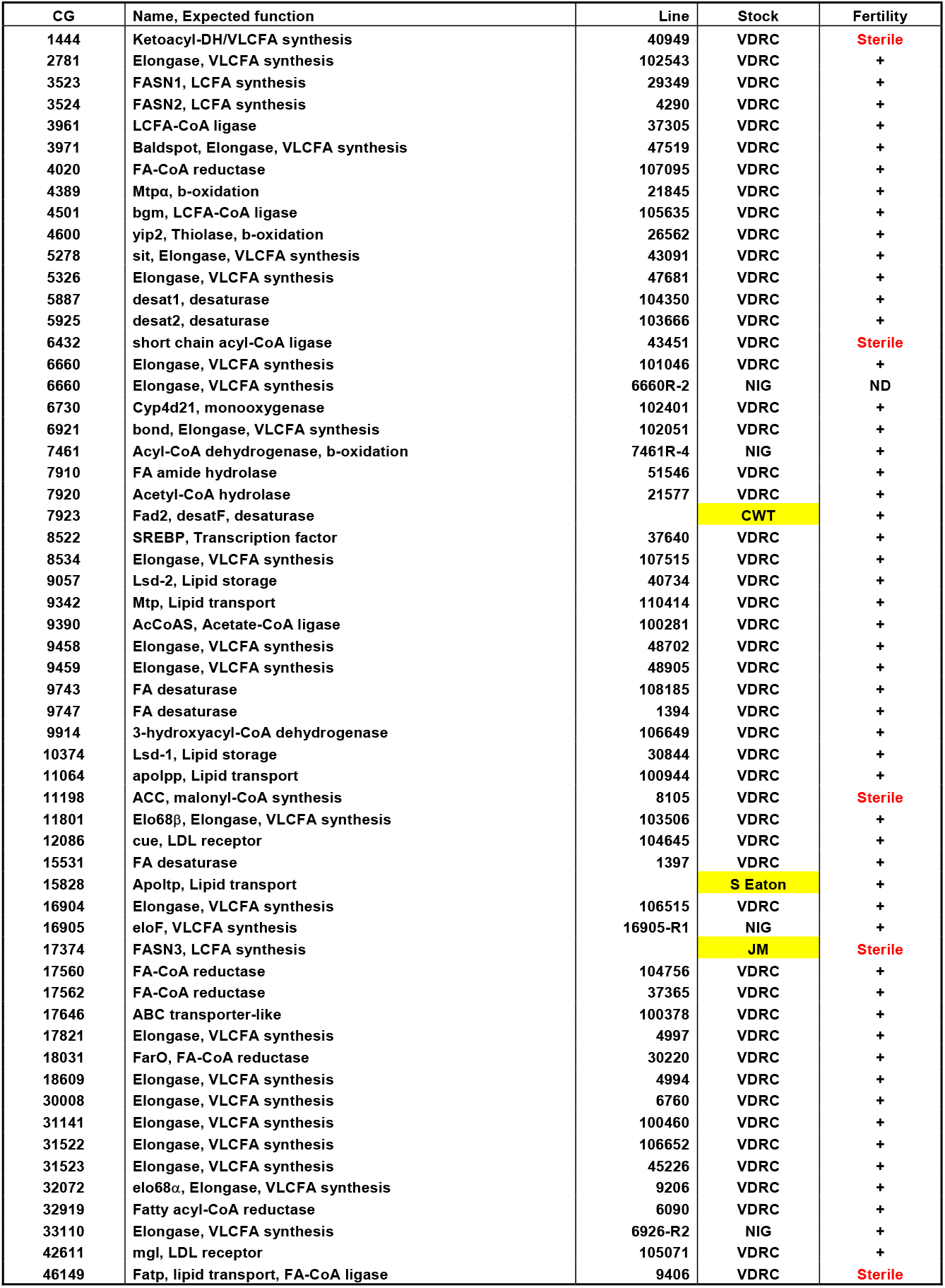
List of the genes screened for female sterility. (fertility column) using the *1407-gal4* driver. RNAi lines were provided by NIG or VDRC; three of them have been generated in S Eaton, CWT or JM laboratories (Stock column). *UAS-RNAi* lines that produced a sterile phenotype were re-used with the *promE-gal4* driver, except 6660R-2 used to knockdown *elo*^*CG6660*^.

**Table EV2.** Statistical and CHC detailed analyses: Analysis of *1407>CG6432-Ri* (*6432i*) 4-5-day old males (top) or females (bottom). First column: CHC identities; elemental composition is indicated as the carbon chain length followed by the number of double bonds; Me-are mbCHCs. CHCs are expressed in ng/ fly (Tot) or in percentages relative to total CHC amount as the mean (± SEM) of CHCs produced by 10 flies kept for 4 days at 25°C.

| **Males CHC** | ***P*** | ***6432i-C*** | ***6432i*** |
| --- | --- | --- | --- |
| Tot | 0.24 | 1634.1 ± 91.8 | 1840.3 ± 146.8 |
| 9-T | **0.003** | 2.50 ± 0.25 | 3.78 ± 0.28 |
| 7-T | **0.002** | 51.33 ± 1.05.2 | 55.7 ± 0.61 |
| 5-T | 0.25 | 3.72 ± 0.18 | 4.14 ± 0.31 |
| 23 :0 | **0.004** | 13.54 ± 0.41 | 14.69 ± 0.30 |
| Me-24 | **<.001** | 3.03 ± 0.24 | 0.38 ± 0.06 |
| 9-P | **0.003** | 1.27 ± 0.14 | 2.24 ± 0.14 |
| 7-P | **<.001** | 5.62 ± 0.19 | 10.24 ± 0.53 |
| 5-P | 0.13 | 0.12 ± 0.07 | 0.34 ± 0.11 |
| 25 :0 | 0.50 | 1.85 ± 0.19 | 2.05 ± 0.22 |
| Me-26 | **<.001** | 6.68 ± 0.43 | 0.42 ± 0.11 |
| 27 :0 | **<.001** | 1.20 ± 0.15 | 2.37 ± 0.21 |
| Me-28 | **<.001** | 7.76 ± 0.61 | 0.62 ± 0.10 |
| 29 :0 | **<.001** | 0.47 ± 0.11 | 2.66 ± 0.24 |

| **Females CHC** | ***P*** | ***6432i-C*** | ***6432i*** |
| --- | --- | --- | --- |
| Tot 23-29 | 0.26 | 1291.6 ± 64.7 | 1574.7 ± 236.1 |
| 9-T | 0.25 | 0.70 ± 0.11 | 0.89 ± 0.11 |
| 7-T | 0.75 | 3.32 ± 0.38 | 3.47 ± 0.29 |
| 5-T | **0.03** | 0.26 ± 0.06 | 0.60 ± 0.13 |
| 23 :0 | 0.05 | 11.22 ± 0.62 | 13.39 ± 0.84 |
| 7,11-PD | **0.04** | 2.44 ± 0.30 | 1.68 ± 0.16 |
| Me-24 | **<.001** | 1.88 ± 0.23 | 0.09 ± 0.01 |
| 9-P | 1 | 3.14 ± 0.31 | 3.14 ± 0.37 |
| 7-P | 0.96 | 2.07 ± 0.18 | 2.06 ± 0.19 |
| 5-P | 0.42 | 0.16 ± 0.06 | 0.23 ± 0.06 |
| 25 :0 | 0.29 | 5.24 ± 0.28 | 5.83 ± 0.48 |
| 7,11-HD | **<.001** | 18.84 ± 0.72 | 13.84 ± 0.56 |
| Me-26 | **<.001** | 16.37 ± 0.57 | 0.60 ± 0.14 |
| 9-H | 0.05 | 3.12 ± 0.14 | 2.56 ± 0.23 |
| 7-H | **0.01** | 2.20 ± 0.16 | 2.90 ± 0.18 |
| 5-H | **0.01** | 0.05 ± 0.02 | 0.19 ± 0.05 |
| 27 :0 | **<.001** | 2.78 ± 0.23 | 10.17 ± 0.44 |
| 7,11-ND | **<.001** | 14.82 ± 0.52 | 28.95 ± 1.55 |
| Me-28 | **<.001** | 9.23 ± 0.42 | 1.37 ± 0.34 |
| 9-N | 0.11 | 0.19 ± 0.07 | 0.40 ± 0.10 |
| 7-N | 0.06 | 0.22 ± 0.08 | 0.44 ± 0.08 |
| 29 :0 | **<.001** | 0.65 ± 0.07 | 6.50 ± 0.55 |

